# PCAT: An integrated portal for genomic and preclinical testing data of pediatric cancer patient-derived xenograft models

**DOI:** 10.1101/2020.06.22.165282

**Authors:** Juechen Yang, Qilin Li, Nighat Noureen, Yanbing Fang, Raushan Kurmasheva, Peter Houghton, Xiaojing Wang, Siyuan Zheng

**Author notes:** These authors contributed equally (Juechen Yang, Qilin Li). Corresponding author: S.Z. or X.W.

## Abstract

Although cancer is the leading cause of disease-related mortality in children, the relative rarity of pediatric cancers poses a significant challenge for developing novel therapeutics to further improve prognosis. Patient-derived xenograft (PDX) models, which are usually developed from high-risk tumors, are a useful platform to study molecular driver events, identify biomarkers, and prioritize therapeutic agents. Here we develop <u>P</u>DX for <u>C</u>hildhood <u>C</u>ancer <u>T</u>herapeutics (PCAT), a new integrated portal for pediatric cancer PDX models. Distinct from previously reported PDX portals, PCAT is focused on pediatric cancer models and provides intuitive interfaces for querying and data mining. The current release comprises 324 models and their associated clinical and genomic data, including gene expression, mutation, and copy number alteration. Importantly, PCAT curates preclinical testing results for 68 models and 79 therapeutic agents manually collected from individual agent testing studies published since 2008. To facilitate comparisons of patterns between patient tumors and PDX models, PCAT curates clinical and molecular data of patient tumors from the TARGET project. In addition, PCAT provides access to gene fusions identified in nearly 1,000 TARGET samples. PCAT was built using R-shiny and MySQL. The portal can be accessed at http://pcat.zhenglab.info or http://www.pedtranscriptome.org.

## INTRODUCTION

Cancer is the leading cause of disease-related mortality in children. Approximately 300,000 children under age of 14 are diagnosed with cancer globally each year (1). In 2019, about 11,000 new diagnoses were reported in the US, with about 1,200 disease-caused deaths (2). Over the last five decades, intensive treatments combining surgical resection, radio- and chemotherapy have significantly improved the outcomes of pediatric cancer. For Instance, five-year survival rate has increased from 58% in mid 1970s to 83% in 2014, with the mortality rates declined by 65% from 1970 to 2016 (2). However, prognosis for relapse patients remains poor, and intensive treatments cause long-term health problems such as secondary cancers, cardiovascular diseases, cognitive disabilities for brain tumors, etc. (3). Many pediatric cancers such as Ewing sarcoma lack targeted therapy. Therefore, continuous efforts on finding new therapeutic targets and developing less-toxic treatments for children with cancer are important for further improving prognosis and mitigating long-term health problems for survivors.

Pediatric cancers constitute about 1% of annual new cancer diagnoses. This small population can be further split into many disease entities thus each has only a very small number of cases. This rarity poses a significant challenge for translational research, as collecting and testing agents in patients, especially those of ultra-rare subtypes, is difficult. Patient-Derived Xenograft (PDX) models have been used for the past four decades to alleviate these difficulties. These models are generated by implanting patient tumors into immune-deficient rodents and have been shown to retain histological and genomic features of the original tumors (4,5). Preclinical testing of these models to therapeutic agents has generated highly valuable insights to guide clinical trials in patients. Moreover, advances in sequencing and other high throughput technologies now allow comprehensive molecular characterization of these models (6). The resulting genomic profiles provide a repertoire for guiding development of targeted therapies, identifying biomarkers for drug sensitivity, and for understanding the genetic basis of resistance.

Here, we introduce *P*DX for *C*hildhood c*A*ncer *T*herapeutics (PCAT), a new database of pediatric cancer PDX models. PCAT currently stores information of 324 PDX models spanning all major cancer types seen in children, including some very rare subtypes. Of these models, 309 have at least one type of genomic profiling data (somatic mutation, n=289; expression/fusion, n=244; copy number, n=282). Preclinical testing data are available for 68 models across 79 therapeutic agents. To facilitate comparisons of PDXs and patient tumors, PCAT curated clinical and molecular data from TARGET so that patterns learned from PDXs can be easily replicated in patient tumors. User friendly interfaces were constructed for searching and data mining. PCAT is freely available without the need for registration at http://pcat.zhenglab.info or http://www.pedtranscriptome.org.

## DATA CONTENT

In the current release, PCAT hosts information of 324 pediatric cancer PDX models. These models reflect cancers commonly observed in children, including acute lymphocytic leukemia (ALL, n=95), osteosarcoma (OS, n=45), neuroblastoma (NBL, n=40), medulloblastoma (MBL, n=25), rhabdomyosarcoma (RMS, n=20), Wilms tumor (WT, n=14), atypical teratoid rhabdoid tumors (ATRT, n=12), glioblastoma (GBM, n=12), ependymoma (n=12), Ewing sarcoma (ES, n=11), and 36 others (Figure 1A). In addition to diagnosis, demographic information is available for 294 models. The average age of the tissue donors was 8 years old. Thirty-seven percent of the models were derived from relapsed, post-treatment, or progressing diseases, whereas 63% were derived from tumors at diagnosis.

**Figure 1.**
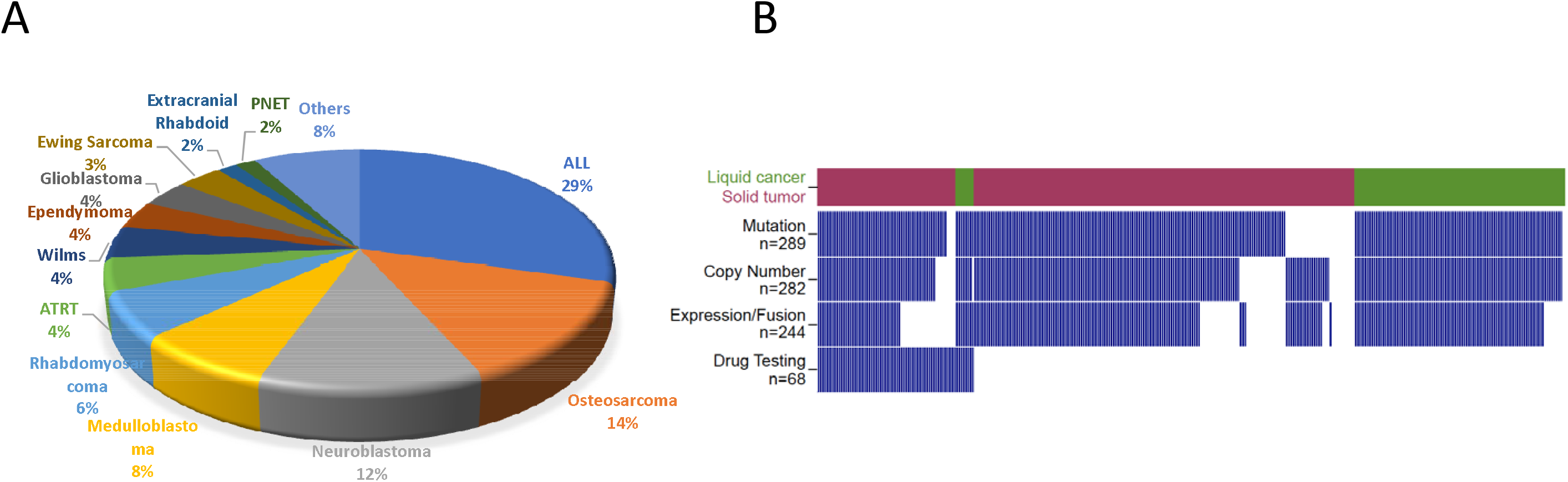
A summary of PDX models in the current PCAT release. (A) A pie chart illustrating histology of the 324 models. (B) An overview of molecular and preclinical testing data of the 324 models. Each column represents one model. Tumors are separated into liquid cancer (dark green) and solid tumors (maroon). Blue bar denotes data is available for the model.

Mutation and copy number data were curated for 289 models either from PPTP (Pediatric Preclinical Testing Program) (7) or PPTC (Pediatric Preclinical Testing Consortium) (8). Gene level copy number changes were obtained by discretizing copy number values into homozygous deletion (−2), heterozygous deletion (−1), neutral (0), gain (1), and amplification (2) using GISTIC2 (9). RNA sequencing based gene expression and fusions were obtained from PPTC (8). Preclinical testing results of 68 models over 79 therapeutic agents were manually collected from individual agent testing studies published since 2008. Drug responses are categorized into six levels, progressive disease 1 (PD1, value 1); progressive disease 2 (PD2, value 2); stable disease (SD, value 3); partial response (PR, value 4); complete response (CR, value 5); maintained complete response (MCR, value 6). Detailed explanation of these six drug response levels for solid tumors and blood cancers can be found on the ‘help’ page of the website. A summary of PDX molecular data is shown in Figure 1B. A different visualization (UpSet plot) is provided in Supplementary Figure 1.

In addition to PDX models, PCAT also curated patient tumor data from TARGET (Supplementary Figure 2). These data enable users to examine patterns observed from PDXs in patient tumors, and to further perform analyses that are not feasible using models such as survival analysis. Clinical, mutation, and expression data of the TARGET dataset were downloaded from GDC data portal (Supplementary Table 1). Copy number segmentation files were downloaded from TARGET Data Matrix and were further analyzed by GISTIC2 to ensure compatibility with those of PDXs (9). Only high confidence mutations called by at least two callers (MuSE (10), MuTect2 (11), SomaticSniper (12) and VarScan2 (13)) were included in our database.

**Figure 2.**
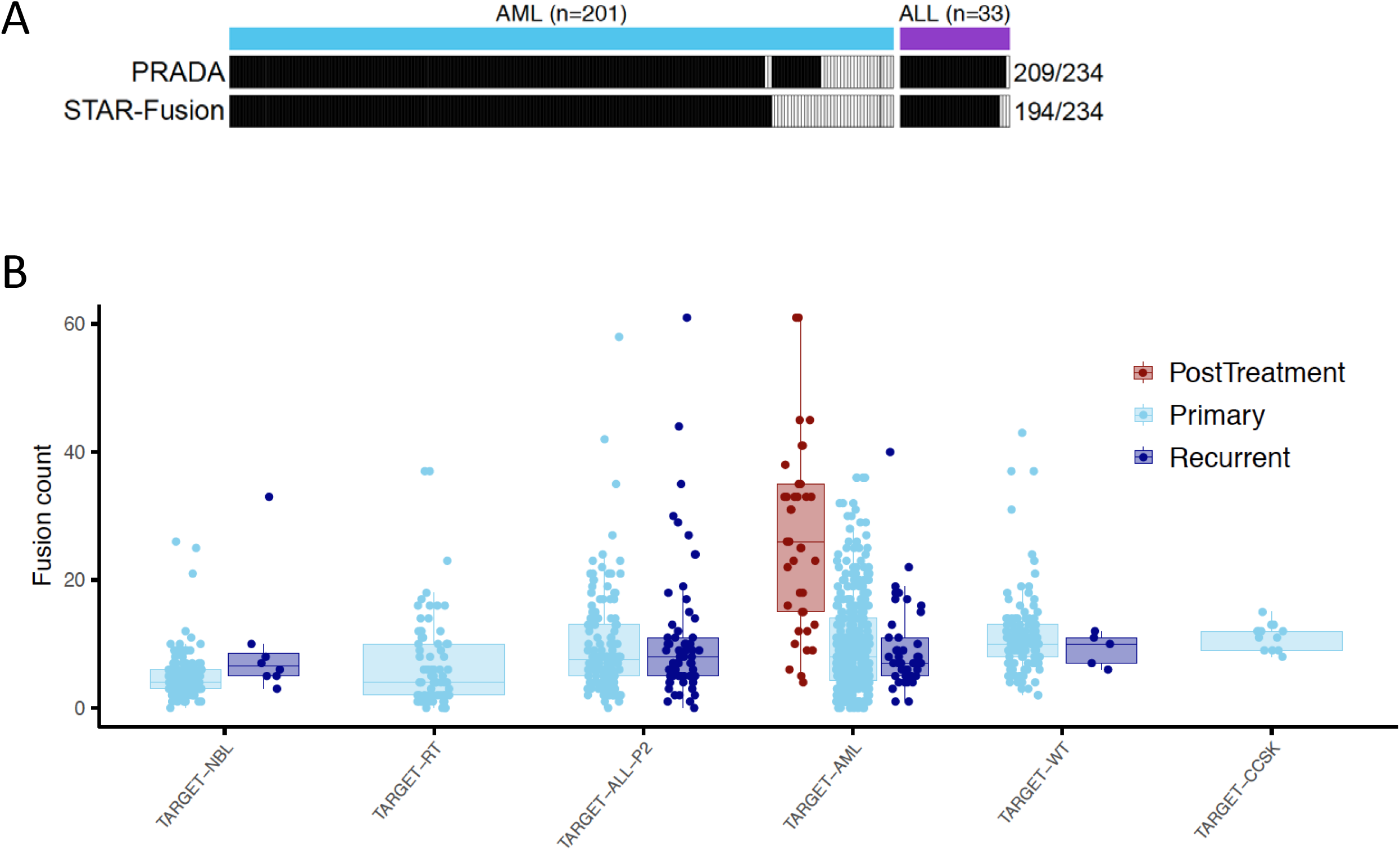
Gene fusions identified in TARGET samples. (A) Benchmarking of fusion identification against 234 driver events annotated in ALL and AML clinical data. Each column represents one fusion. Solid color indicates the fusion was found by one of the tools. (B) Distribution of fusion events across cancer types and sample types (NBL, neuroblastoma; RT, rhabdoid tumor; ALL, acute lymphoblastic leukemia; AML, acute myeloid leukemia; WT, Wilms tumor; CCSK, clear cell sarcoma kidney). Each dot represents a cancer sample. Fusion counts of post-treatment samples are significantly higher than primary and recurrent samples in ALM (both p values < 0.001, Wilcoxon rank sum test).

Gene fusions are a very important group of cancer drivers, particularly for childhood cancer. To catalogue gene fusions as a community resource, we employed our in-house fusion caller PRADA (14) and a well benchmarked tool STAR-Fusion (15) on 943 TARGET samples (Supplementary Table 2). This analysis identified a total of 8912 fusions by the two callers, with 3718 by PRADA, 5980 by STAR-fusion, and 786 called by both callers. We benchmarked our fusion identification using driver fusion events annotated for some ALL and AML patients in the clinical data. Of the 234 driver fusion events, PRADA identified 209 (89%) and STAR-Fusion identified 194 (83%). Combined, these two tools identified 90% of the total 234 fusions (Figure 2A). We next broke down these fusions by cancer type and sample type. As expected, considerable heterogeneity was observed in fusion loads in each cancer type (Figure 2B). Interestingly, post-treatment AML samples demonstrated significantly higher number of fusions than primary and recurrent samples (both p values < 0.001, Wilcoxon rank sum test), consistent with the anticipation that cytotoxic chemotherapy causes DNA breaks leading to increased fusion rates. Few post-treatment samples were available for other cancer types thus were excluded from this analysis.

## WEB INTERFACE AND DATA DOWNLOAD

PCAT web interface is organized into a resource and two analysis sections. Each analysis section consists of several functional modules. The resource section is the interface to the major data stored on PCAT, including the 324 PDXs and gene fusions. When searching for PDXs, users can specify histology, mutation, or gene fusion as search criteria. Results will be returned in a tabular format divided into clinical information, mutation, fusion, and preclinical testing if available. An example of the PDX summary page is shown in Figure 3A. Similarly, fusion search results will be returned also in a tabular format listing fusion and the case ID where this fusion is found. Clicking each fusion links to a page that summarizes the prevalence of the fusion in disease cohorts. The fusion detail page displays technical parameters of the fusion identification, a circos plot illustrating all fusions identified in the sample, and the expression of the two partner genes (Figure 3B).

**Figure 3.**
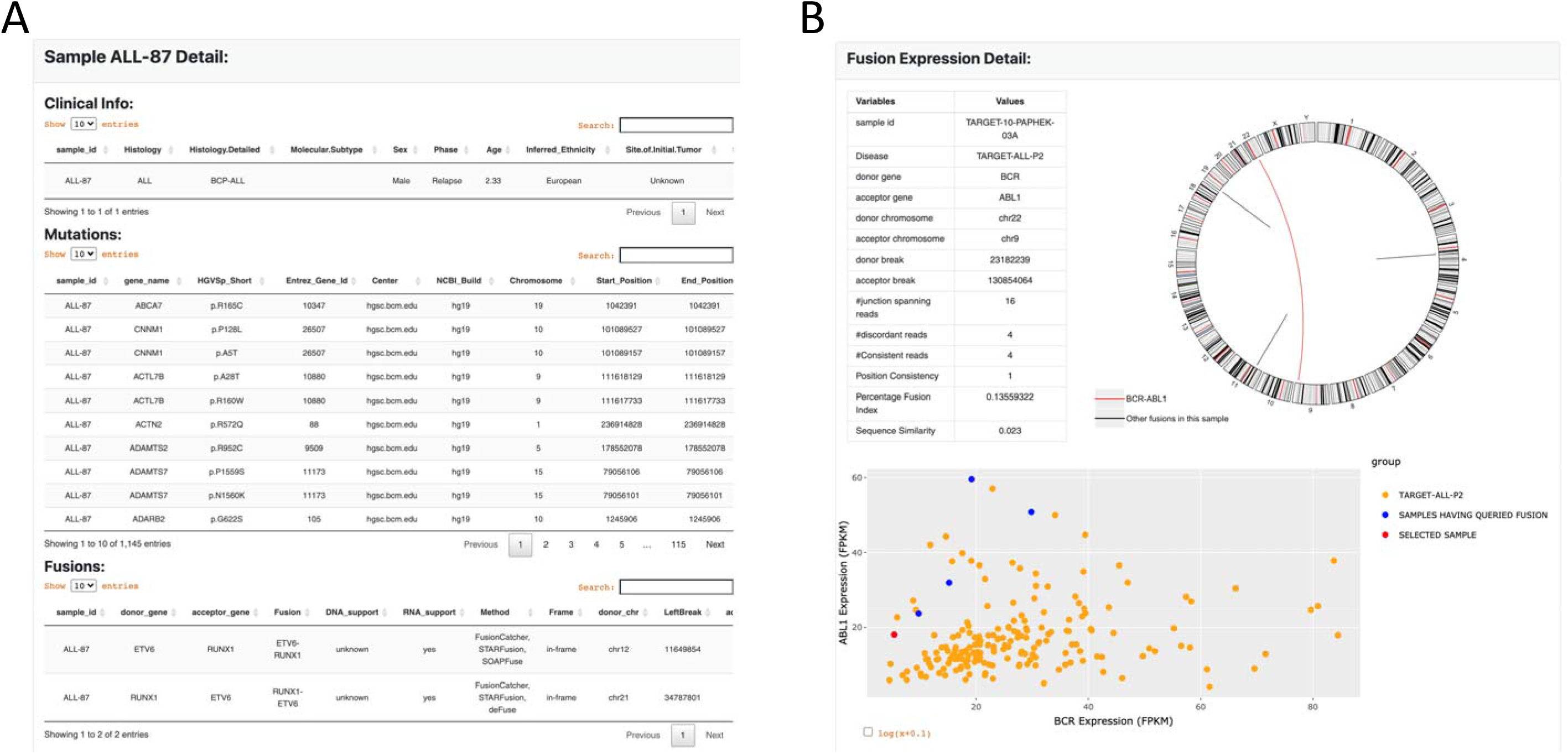
The display page of PDX and fusion search results. (A) PDX Information is displayed in a tabular page divided into clinical, mutation, fusion, and preclinical testing data if available. (B) Fusion page displays identification information (evidence, junctions, etc), a circos plot illustrating all fusions identified in this case, and a scatter plot showing expression of the two partner genes. BCR-ABL1 is used in this example (red in circos plot).

The analysis modules are designed to enable exploration of genomic and preclinical data of PDXs and patient tumors. In the current release, PCAT is focused on gene expression data analysis because childhood cancers harbor far fewer mutations than their adult counterparts according to recent large-scale genomic studies (16,17). “Single gene analysis” modules allow users to correlate expression of the input gene with histopathological parameters, patient prognosis, and genomic alterations (copy number and mutation). It also allows users to find genes that share co-expression patterns with the input gene in a selected dataset. “Multiple gene analysis” modules allow users to visualize expression patterns of the input genes (“visualization” module). The “single sample gene set enrichment (ssGSEA) analysis” module allows users to aggregate the expression of an input gene set into a single score. The co-expression module returns pairwise expression correlation based on the select dataset. If users choose to remove lineage effect, PCAT will z-score transform the expression data for each tissue of cancer origin before calculating expression correlation. Finally, PCAT allows correlation of gene expression and mutation with preclinical testing results in the “preclinical testing” module.

For all modules, PCAT provides download links for returned results so that users have the opportunity to reproduce and customize figures for publications and other purposes. Fusion results for TARGET can be found in Supplementary Table 2.

## ANALYSIS MODULES

We use examples to demonstrate the utility of the analysis modules in generating and testing hypotheses. First, we show expression of PDGFRA, a marker of mesenchymal stem cells, in PDX models (Figure 4A). As expected, PDGFRA is highly expressed in cancers of mesenchymal origin including extracranial rhabdoid cancer and osteosarcoma. This pattern is replicated across TARGET cohorts (Supplementary Figure 3). Next, we show the correlation between CDKN2A deletion and expression. CDKN2A is a well-established tumor suppressor and is frequently deleted in cancer. In ALL models, CDKN2A deletion is strongly associated with decreased expression (Figure 4B). The same pattern is observed in osteosarcoma (data not shown), a cancer type with high genomic instability (18). These data collectively show DNA deletion is a common mechanism to inactivate CDKN2A in cancer.

**Figure 4.**
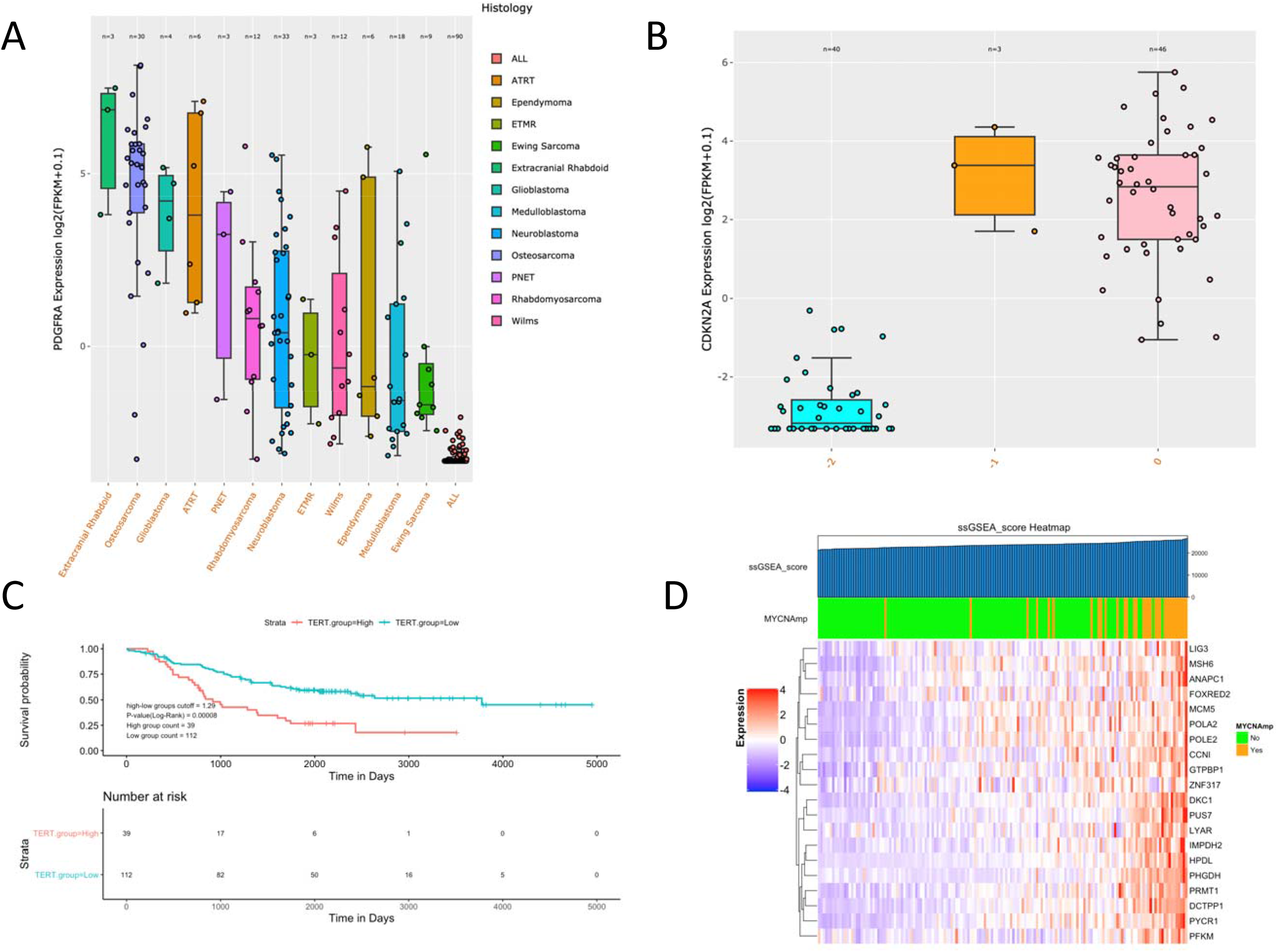
Examples demonstrating the utility of the analysis modules. (A) Expression of PDGFRA, a mesenchymal stem cell marker, across PDX models. (B) Correlation of CDKN2A expression and deletion in ALL PDXs. “−2”, homozygous deletion; “−1”, shallow deletion; “0” copy number neutral. (C) Correlation of TERT expression and patient overall survival in TARGET neuroblastoma. TERT expression is divided into high and low based on median expression. (D) ssGESA results for top 20 MYCN targets in TARGET neuroblastoma. The color bar on top of the heatmap indicates MYCN status per clinical data annotation. Samples are ordered ascendingly per ssGSEA score, as indicated by the top bar.

The “survival analysis module” allows users to perform survival analysis using both clinical parameters and gene expression. PCAT supports univariate and multivariate survival analysis by deploying the R package survminer. When no query gene is input, users can perform survival analysis using clinical parameters. To correlate gene expression with clinical outcome, PCAT provides four methods to divide gene expression into groups, including auto-calculated threshold, mean value, median value, and customized cutoff. The auto-calculated threshold is calculated by testing all possible cut-off values between the top 20% and bottom 20% samples based on the expression of the gene and adopts the value that best separates the clinical outcomes of the high and low groups. Kaplan - Meier plot is generated for visualization. Users may also choose one or several clinical parameters as co-variates to conduct multivariate survival analysis. A forest plot will be generated to show the hazard ratio of each variate generated in the analysis. To demonstrate the utility of this module, we use TERT and neuroblastoma as an example. High TERT expression, an indicator of active telomerase, predicts high risk tumors in neuroblastoma (19). Using either mean or median to split the cohort, PCAT shows high TERT expression is significantly associated with worse overall survival in the TARGET neuroblastoma dataset (Figure 4C). This correlation holds even when MYCN amplification is added as a covariate to the analysis (Supplementary Figure 4), suggesting TERT expression is an independent prognostic factor.

We use a list of 20 MYCN targets identified by shRNA screening (20) to demonstrate utility of the “ssGSEA” module. We first ran ssGSEA using these genes in PDXs. The output clearly showed higher scores in neuroblastoma models than others, suggesting this group of genes is upregulated in neuroblastoma (Supplementary Figure 5). We then re-ran ssGSEA using the TARGET neuroblastoma dataset. We observed that MYCN amplification was strongly enriched in samples with higher ssGSEA scores (hence higher expression of the target genes) (Figure 4D), verifying the positive regulation of these genes by MYCN.

**Figure 5.**
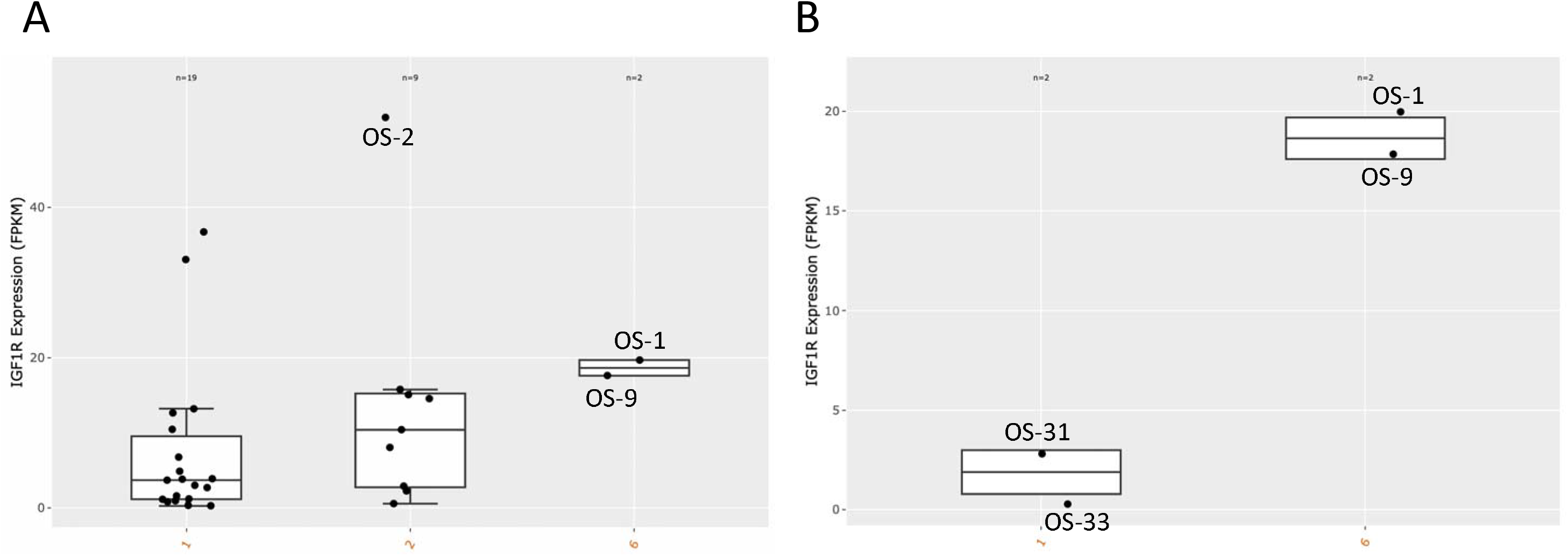
Molecular correlates of drug responses. (A) Model response to 19D12, an IGF1R antibody inhibitor, is correlated with high expression of the target gene IGF1R. X axis reflects the RECIST criteria: 1-2, progressive disease; 6, maintained complete response. (B) Limiting the analysis to osteosarcoma models only sustains the association, suggesting it is not a tissue effect i.e. osteosarcoma has high IGF1R expression and the drug works better on osteosarcoma. In both figures, a minimum of two samples was required per each response group.

Finally, we use 19D12 (also known as SCH717454) inhibitor to demonstrate the utility of the preclinical testing module. SCH717454 is a fully human antibody inhibiting the insulin-like growth factor 1 receptor (IGF1R) (21). Correlating IGF1R expression and response to 19D12 reveals that maintained complete response (MCR) demonstrated by two osteosarcoma models (OS-1, OS-9) is associated with high IGF1R expression (Figure 5A). The same pattern was observed when limiting this analysis to osteosarcoma models only (Figure 5B). OS-2, another osteosarcoma model with much higher IGF1R expression, shows limited response to 19D12 (PD2). Interestingly, this model has much higher expression of IGF1 and IGF2 than the two sensitive models (Supplementary Figure 6). Despite a small sample size, these observations suggest that high IGF1/2 expression may be an escape mechanism to 19D12 inhibition for osteosarcomas.

## SUMMARY AND FUTURE DIRECTIONS

In this work, we describe PCAT, a new resource for childhood cancer PDX models. Previously published portals such as PDXFinder (22) has a larger repository than PCAT. However, PCAT is distinct in its collection of childhood cancer PDXs. For instance, searching “neuroblastoma”, a malignancy of the peripheral nervous system commonly seen in children, found one model on PDXFinder, but 40 models on PCAT. Among the PCAT functions/features not provided by PDXFinder are the intuitive interface that has been developed to allow users to explore the genomic and preclinical testing data of these models, as well as facilitation of comparisons between patient tumors and PDX models by integrating TARGET datasets into the portal. The gene fusions of nearly 1,000 tumors is a unique resource that allows users to inquire and examine genes and their potential involvement in fusion events.

The next step for PCAT is to expand its PDX collection. More than 100 new models generated by PPTC and CPRIT GCCRI Core (https://gccri.uthscsa.edu/services/pdx-core/) are currently in the pipeline and will be integrated into PCAT. Most of these models were derived from patients of Hispanic ethnicity thus reflecting a unique demographic patient group in south Texas. Meanwhile, more preclinical testing data and protocols of preclinical testing experiments will be gradually added to the portal.

In addition, more functional modules will be added to facilitate data mining and visualization. For instance, new modules will be added to enhance users’ ability to explore mutations. More importantly, we will implement functions that allow users to compare their own samples to our PDX models so that results of PDX preclinical testing can be a reference to predict the query sample’s sensitivity to therapeutic agents. We envision these new modules will greatly enhance the usability and translational relevance of this resource.

## ACKNOWLEDGEMENT

We are grateful to the PPTP/PPTC group for generating many of the PDX models and their genomic and preclinical data. We are also grateful to colleagues at GCCRI for extensive discussions and their expertise in childhood cancer. Work in Zheng laboratory is supported by CPRIT (RR170055). P.H. and R.K. are supported by CRPIT (RP160716) and NCI (UO1CA199297). X.W. is supported by GCCRI pilot project. The results published here are in whole or part based upon data generated by the TARGET (https://ocg.cancer.gov/programs/target) initiative, phs000218. The data used for this analysis are available at https://portal.gdc.cancer.gov/projects.

---

**Supplementary Figure 1.**
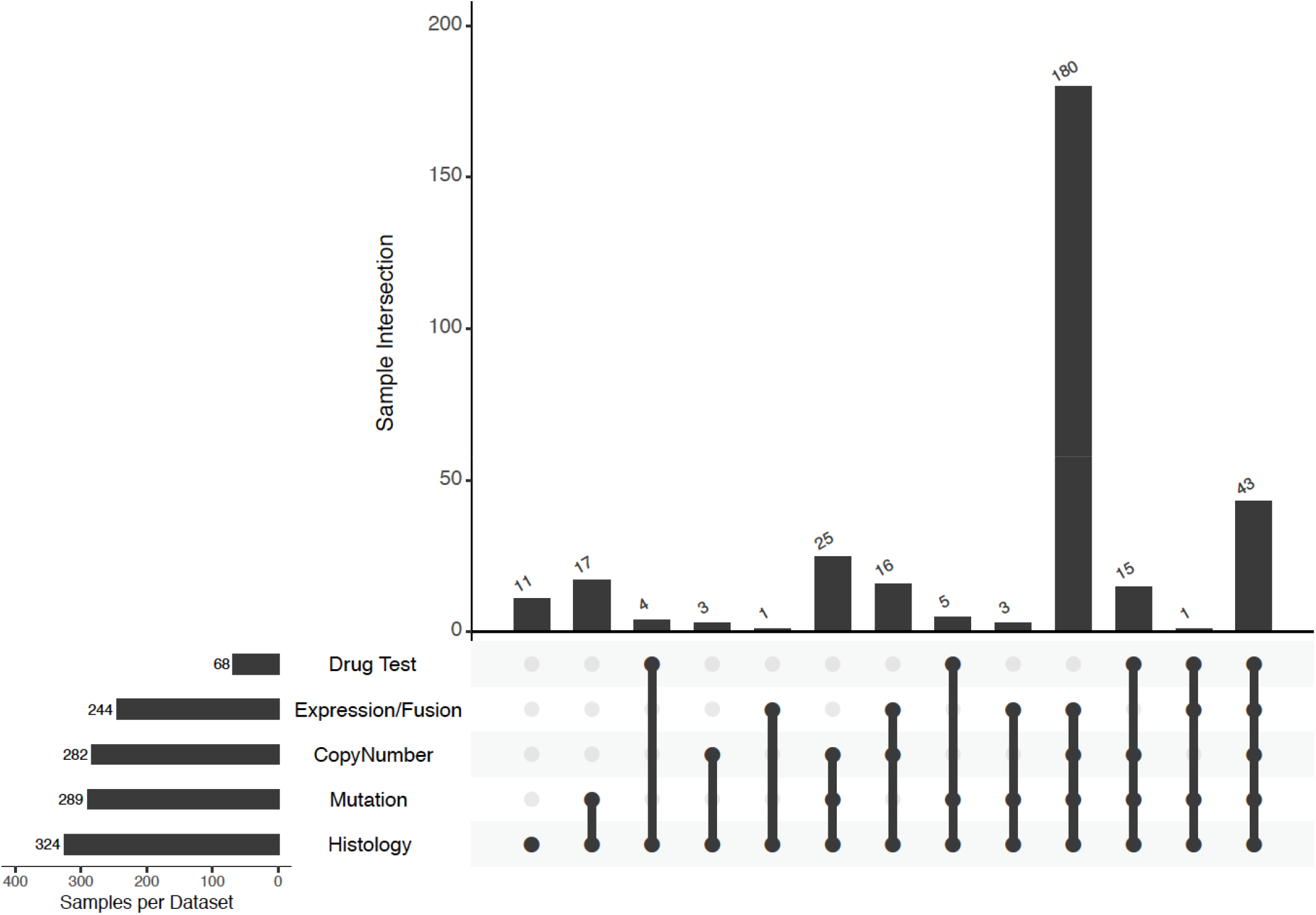
A summary of PDX data currently hosted at PCAT. The bars on the left show the total number of samples for each data type. Bars on top show the number of samples with the overlapping data indicated by the solid connecting dots.

**Supplementary Figure 2.**
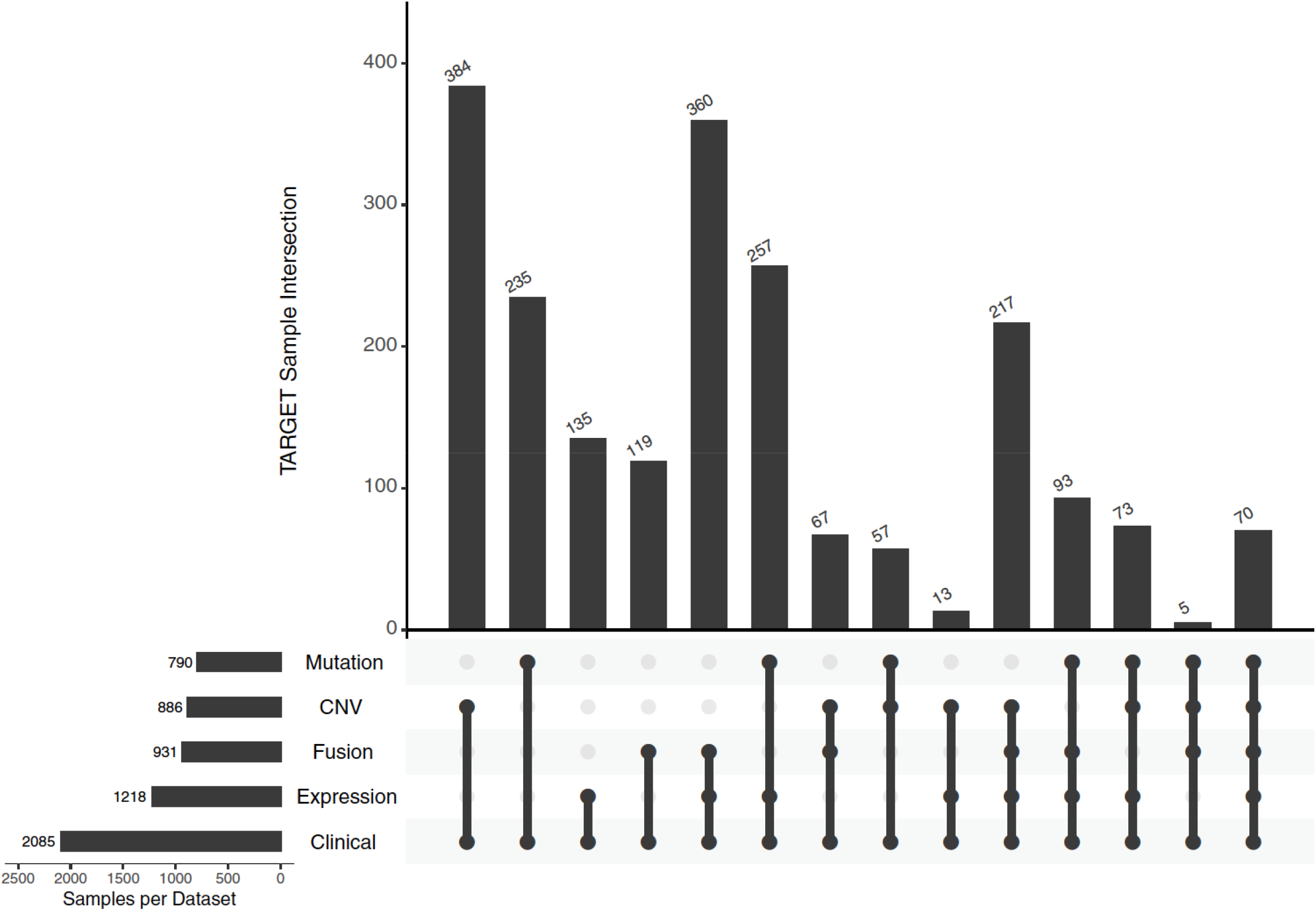
A summary of the TARGET data that have been integrated into PCAT. The bars on the left show the total number of samples for each data type. Bars on top show the number of samples with the overlapping data indicated by the solid connecting dots.

**Supplementary Figure 3.**
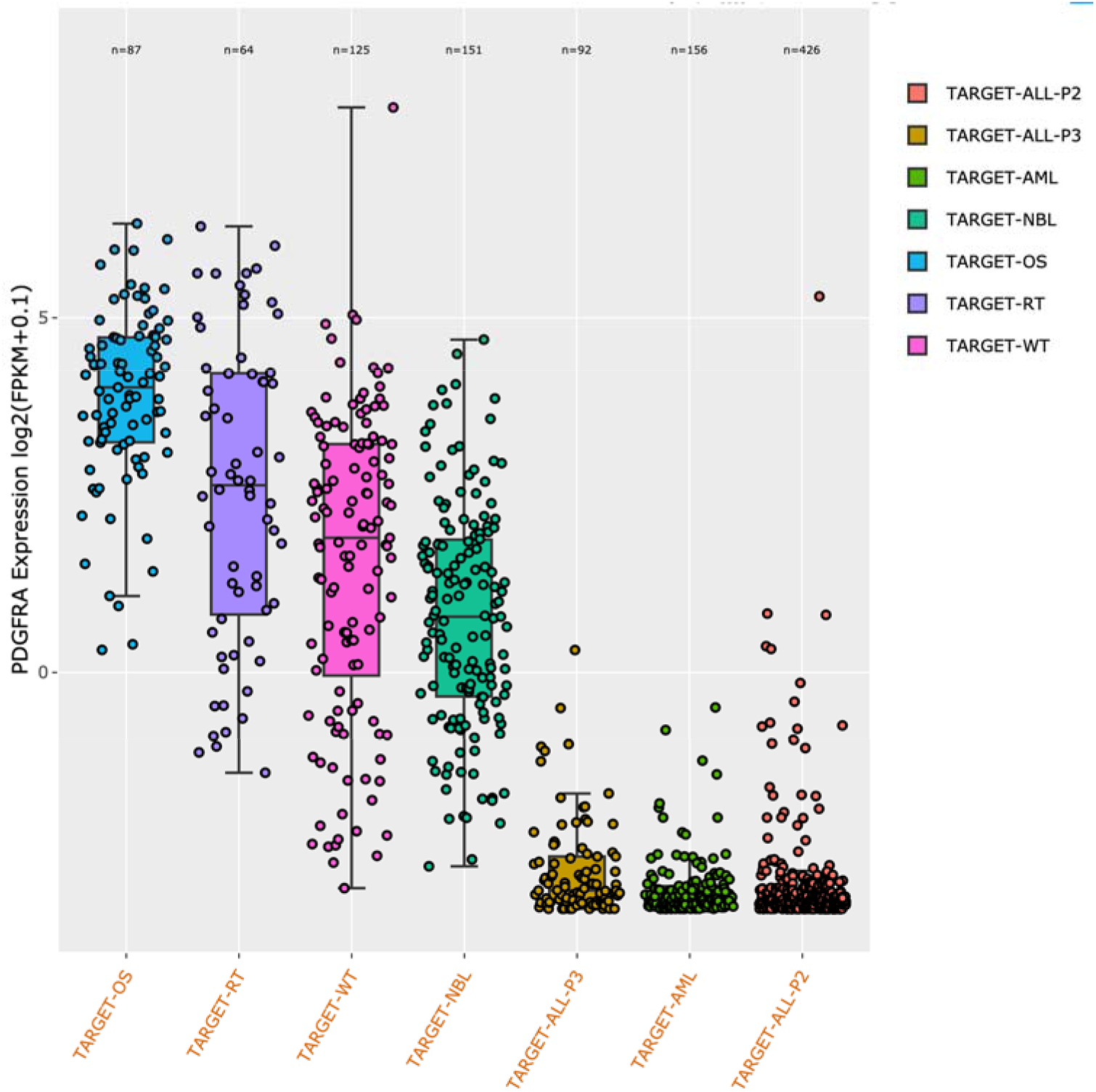
PDGFRA expression across TARGET cohorts. Similar to observations made in PDX models, PDGFRA is highly expressed in cancers of mesenchymal origin including osteosarcoma and rhabdoid cancer but very low in blood cancers.

**Supplementary Figure 4.**
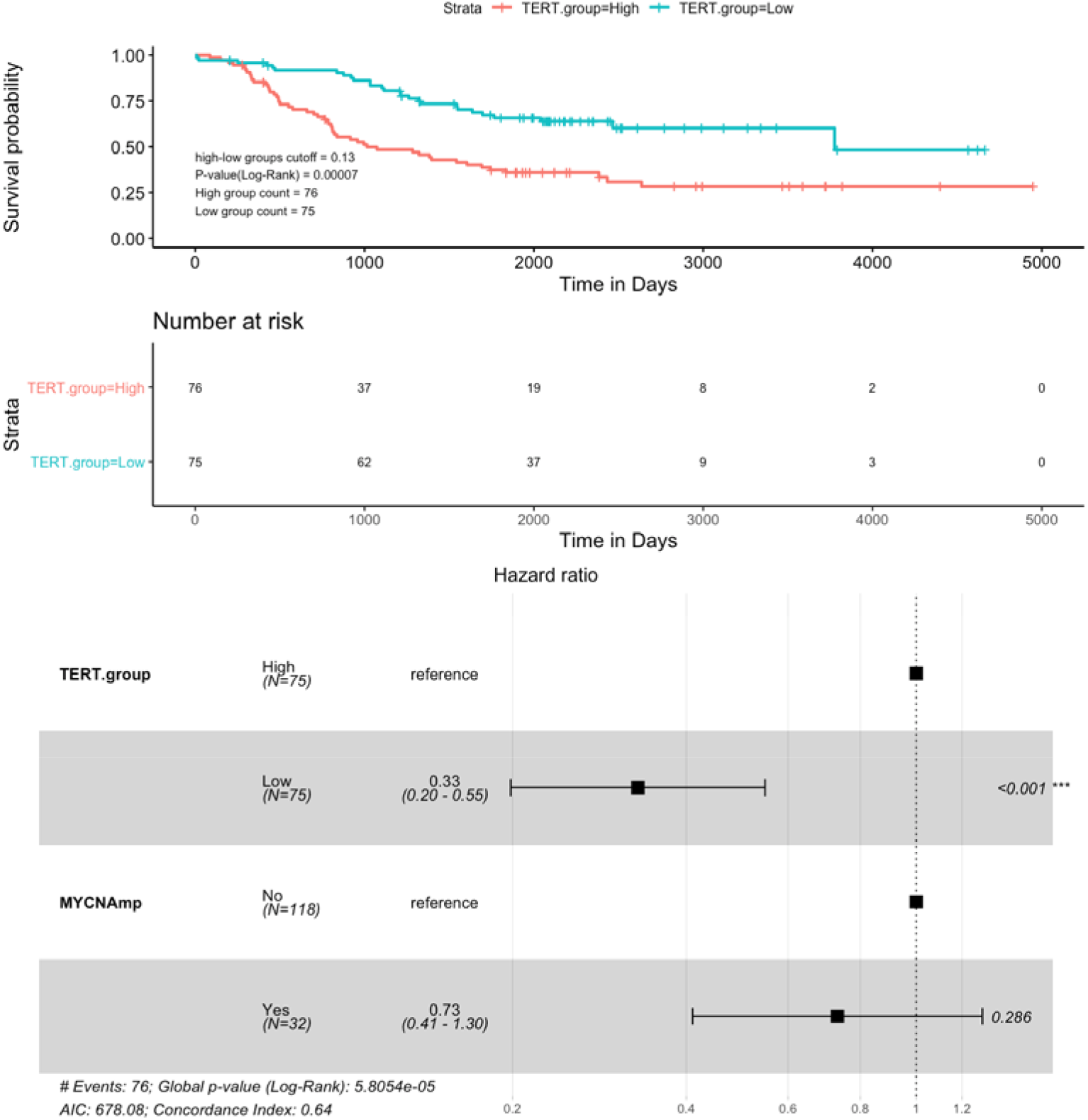
Multivariate survival analysis. This example tests the correlation between TERT expression and neuroblastoma prognosis. MYCN amplification status is included to demonstrate that the prognostic value of TERT expression is independent of MYCN.

**Supplementary Figure 5.**
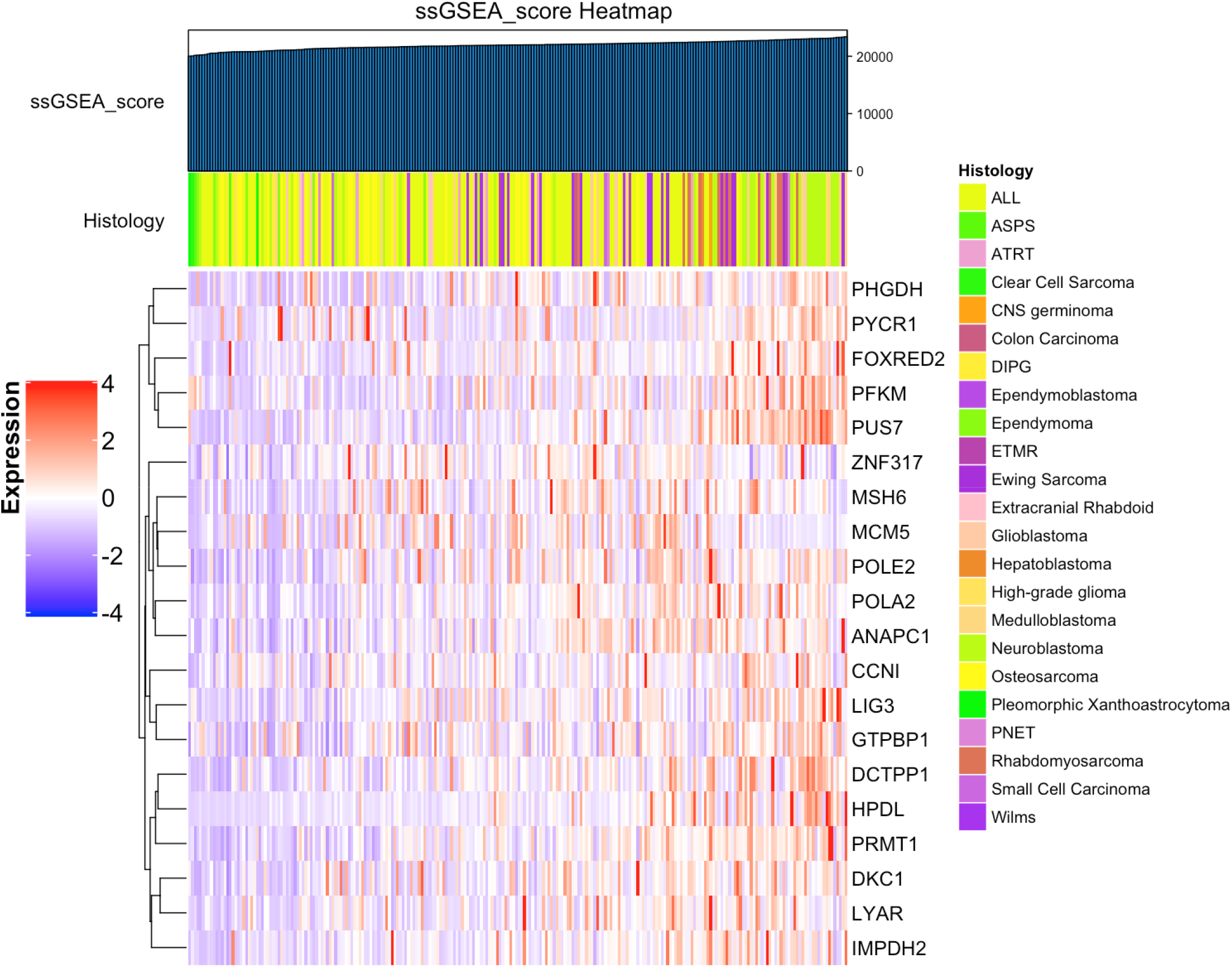
ssGSEA analysis of MYCN targets in PDXs. The top 20 genes positively regulated by MYCN were downloaded from Valentijn et al. PNAS 2012. These genes were identified from shRNA screening in the original study. Each column represents one PDX model. Samples are odered from low to high based on ssGSEA scores.

**Supplementary Figure 6.**
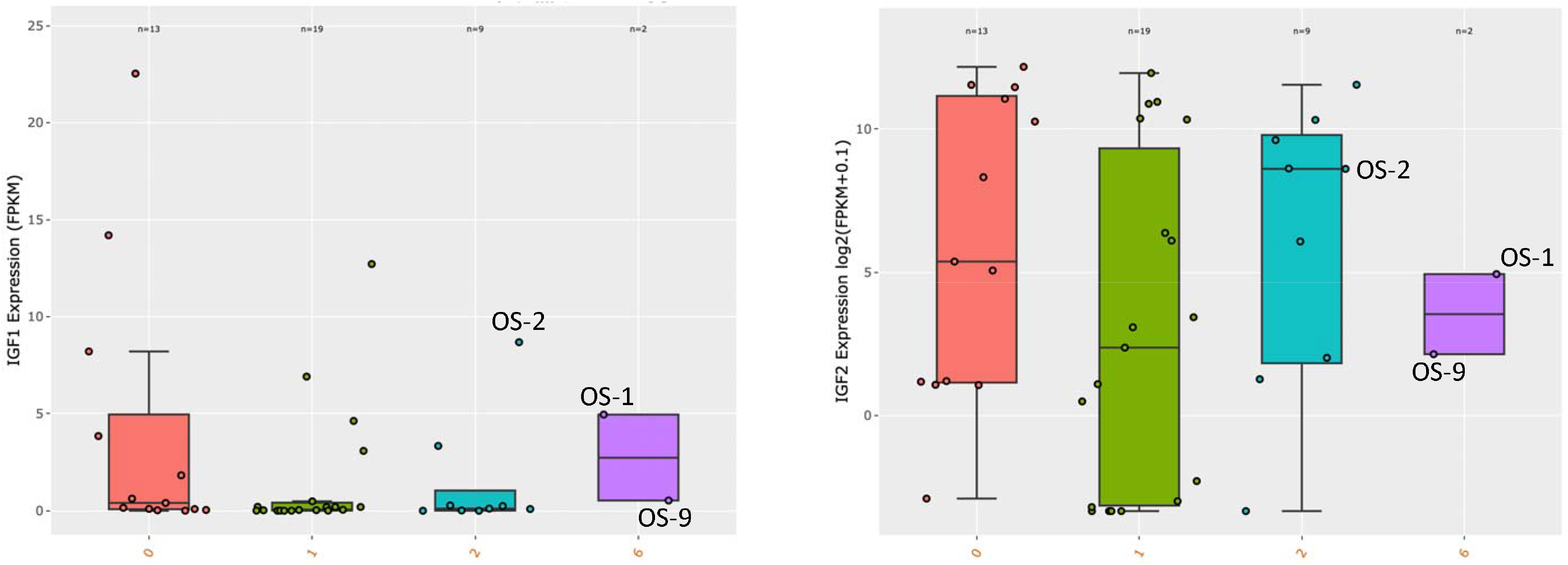
Molecular correlates of drug responses. (A) Although OS-1 and OS-9 show MCR to IGF1R inhibitor 19D12, another osteosarcoma model OS-2 shows no response (progressive disease) despite higher IGF1R expression. This figure shows OS-2 also has higher expression of IGF1 than the two responsive models. X axis reflects the RECIST criteria: 1-2, progressive disease; 6, maintained complete response. (B) OS-2 has higher IGF2 expression than OS-1 and OS-9. Note that in this figure the y axis is in log scale.

